# Glycine detection with a nuclease-stable L-RNA sensor

**DOI:** 10.64898/2026.08.18.745542

**Authors:** Madeline R. Bodin, Xuan Han, Jonathan T. Sczepanski, Ming C. Hammond

## Abstract

Glycine is a vital extracellular signal in bacteria, plants, and the brain. Although RNA-based sensors detect glycine in cells, their extracellular application in native biological environments is limited by enzymatic degradation from nucleases. Mirror-image RNA is nuclease-resistant and preserves the tertiary structure required for RNA function, but synthesizing long L-RNAs such as the 170-nt glycine sensor (glyS) remains challenging. Here, we applied cross-chiral ligation with natural D-RNA ribozymes to assemble a mirror-image L-RNA glycine sensor (L-glyS). Optimization of the ligation conditions enabled up to 68% conversion to the full-length sensor. L-glyS displayed nuclease resistance and maintained glycine-dependent fluorescence in serum, where the original D-glyS lost function. These results establish cross-chiral ligation as a strategy for constructing long, functional L-RNAs and broaden the possible applications of RNA-based sensors to extracellular detection of small molecules.

## INTRODUCTION

Glycine serves as an important extracellular signal across diverse biological systems. In bacteria, glycine is a chemoattractant that guides the cells towards nutrients.^1^ For this reason, plants release glycine as a root exudate for communication and symbiosis with soil bacteria.^2,3^ In mammals, glycine is released from neurons as a neurotransmitter involved in brain development, reflexes, and memory.^4,5^ More recently, glycine release from astrocytes has also been proposed to influence neuronal signaling,^6,7^ motivating tools to study extracellular glycine.

To study glycine dynamics, the RNA-based fluorescent (RBF) sensor glyS is a useful imaging tool.^8^ GlyS incorporates a bacterial riboswitch that selectively binds glycine and undergoes a conformational change to stabilize a light-up RNA aptamer. This sensor visualized drug-induced glycine accumulation in live *E. coli* and enabled the development of the first single-dye ratiometric sensor.^8^ Despite their utility, RBF sensors in general are susceptible to degradation by cellular and environmental ribonucleases (RNases). Thus, imaging extracellular small molecules in their native environments with RBF sensors remains a challenge. To our knowledge, no established platform exists for the continual extracellular expression or display of RNA sensors, and exogenous RNA is readily degraded by RNases.

Chemical RNA modifications, including phosphorothioate linkages and 2′-sugar substitutions,^9,10^ improve RNase resistance but can disrupt the tertiary fold of RNA-based sensors. Although *in vitro* reselection has enabled RNase-resistant fluorogenic aptamers with modified sugars,^11^ this approach can be time-consuming, and the results are bespoke and not readily generalized. An alternative approach that avoids reselection is using mirror-image RNA (L-RNA) to construct the sensor. L-RNA is resistant to RNases, which are stereoselective and therefore do not recognize L-RNA.^12^ L-RNA also preserves the tertiary architecture of RNA aptamers and sensors while maintaining binding to achiral ligands such as guanine^13^ and theophylline.^14^ Critically, both glycine and the Broccoli aptamer fluorophores^15–17^ used by glyS are achiral, suggesting that an L-RNA glyS (L-glyS) would remain functional.

Synthesizing L-glyS is a technical challenge. Solid-phase synthesis with commercial phosphoramidites cannot efficiently produce this 170-nt L-RNA,^18^ and native polymerases cannot incorporate L-nucleotides. Incorporating short D-RNA segments at the termini of L-RNA fragments has enabled ligation by native enzymes,^14^ but this strategy likely will compromise sensor folding. Mirror-image polymerases have been used to generate L-oligonucleotides up to 2.9 kb,^13,19^ but the syntheses of the polymerases themselves are technically demanding and highly specialized.

Instead, we envisioned employing a three-piece cross-chiral ligation strategy^20,21^ in which a D-RNA ligase ribozyme assembles L-RNA fragments into the full-length L-glyS sensor. The D-RNA ligase ribozyme is readily synthesized by standard *in vitro* transcription from a DNA template, and the L-RNA fragments are synthesized using commercial phosphoramidites, which makes this approach more scalable and accessible to other researchers. Another advantage of the solid-phase synthesis and ligation approach over mirror-image polymerases is the ease of incorporating other functional groups, such as fluorophores or affinity tags. Here, we report the synthesis and characterization of an L-RNA-based fluorescent sensor, representing the longest cross-chiral L-RNA assembly to date. The resulting L-glyS clearly outperformed its D-RNA counterpart in RNase-rich serum, retaining glycine-responsive fluorescence. These results establish cross-chiral ligation as a strategy for constructing long, functional L-RNAs for sensing applications.

## RESULTS

### Design Principles for Cross-chiral Ligation of an L-RNA Glycine Sensor

Since cross-chiral ligase ribozymes have previously assembled their own functional mirror image as well as diverse short L-RNAs,^20–23^ we expected that it may be possible to apply this method towards assembling the L-RNA glycine sensor. A three-piece ligation strategy using cross-chiral ligase ribozymes was developed to synthesize L-glyS (Figure S1A). In this approach, a D-RNA ribozyme^20,21^ seamlessly ligates L-RNA fragments in a splint-templated manner.^20–22,24^ Ligation requires that the L-RNA donor fragments bear 5′-triphosphate^20,21^ or 5’-adenosyltriphosphate groups,^23^ and that ligation junctions are paired to complementary L-RNA splints. Previous work showed that cross-chiral ligase ribozymes preferentially ligate 5′-G donor and 3′-C acceptor junctions. These requirements, together with the secondary structure of glyS, guided the selection of ligation sites. Although glyS contains eight potential 5′-GC-3′ junctions (Figure 1A), junctions within the long, stable ghost aptamer stem were excluded to avoid self-hybridization that could compete with splint annealing. To balance fragment lengths with these constraints, ligation junctions were designed to join C75-G76 and C121-G122.

**Figure 1.**
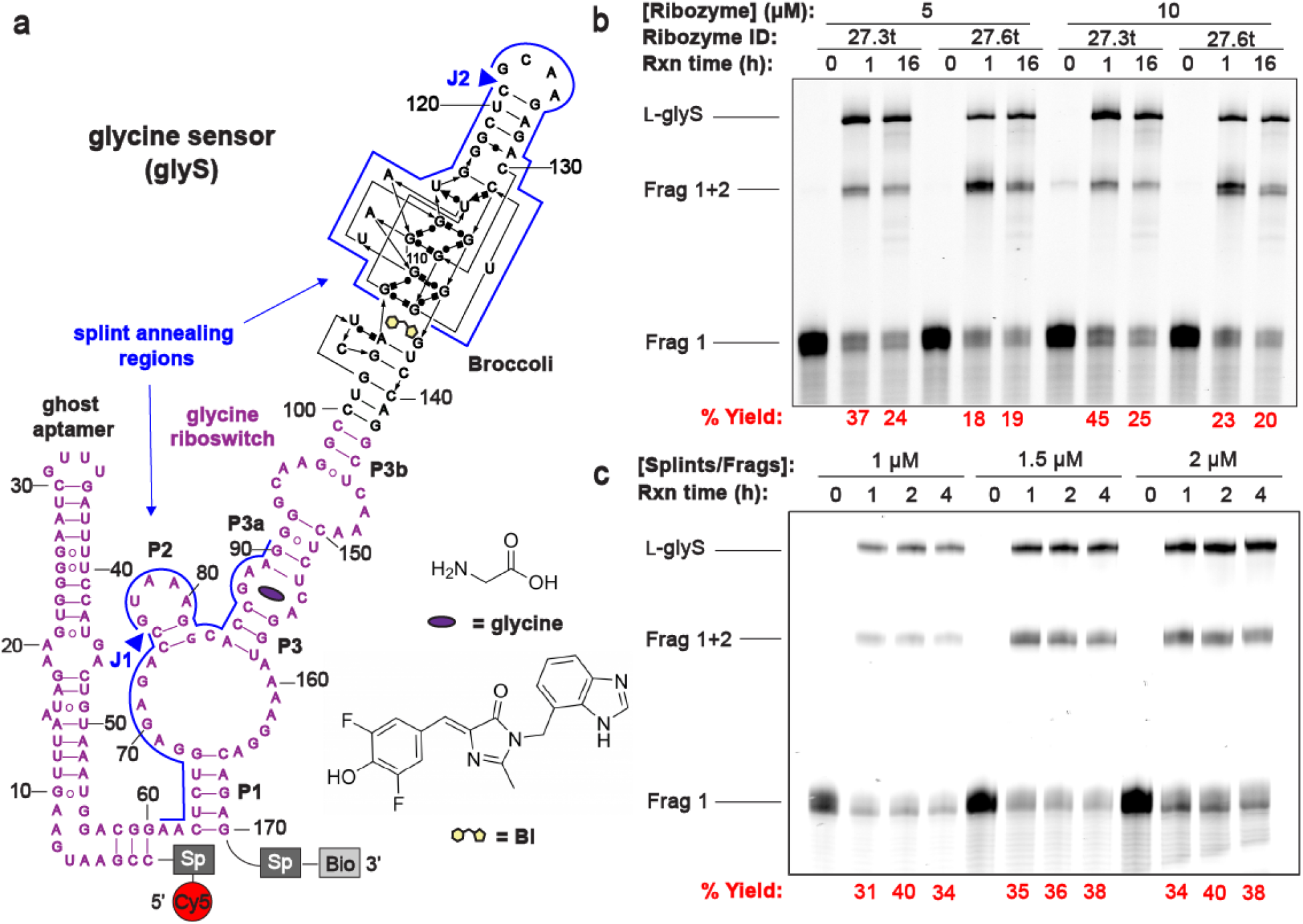
Design and Evaluation of Cross-Chiral Ligations. **(A)** Sequence and secondary structure model of L-RNA glyS sensor. Splint annealing regions are shown with blue lines and ligation junctions (J1, J2) indicated with blue triangles. “Sp” indicates spacer regions, and “Bio” indicates biotin. **(B)** Denaturing PAGE analysis of initial cross-chiral ligations over time with various ribozymes. **(C)** Denaturing PAGE analysis of cross-chiral ligations over time with 5 µM of the 27.3t ribozyme and varying concentrations of L-RNA splints and fragments. Ligations were performed at 23°C with 250 mM NaCl, 250 mM MgCl_2_, and 50 mM Tris (pH 8.5).

The three L-RNA fragments (75, 46, and 49 nt) and two 30-nt L-RNA splints (Table S1) were made by solid-phase synthesis. Fragment 1 contained a 5’-amine group (Figure S1B) for conjugation to NHS-sulfo-Cy5, allowing PAGE visualization and ratiometric imaging. Meanwhile, fragment 3 included a 3′-terminal biotin (Figure S1B) for streptavidin binding to allow future anchoring of L-glyS to biotinylated cell membranes.^25,26^ Importantly, the donor fragments were synthesized with 5’-phosphate groups, which were chemically converted to the 5’-adenosyltriphosphates necessary for cross-chiral ligation activity by incubation with adenosine-5′-diphosphate (ADP)-phosphoimidazolide (ImppA) (Figure S1C).^23^ Mass spectrometry confirmed successful adenosyltriphosphorylation (Figure S2), and PAGE was performed to remove any leftover reactants from the activated L-RNA donor fragments.

### Assembly of the L-RNA Glycine Sensor

To determine whether cross-chiral ligation could produce the full-length L-glyS, multiple ligase ribozymes and reaction conditions were evaluated. The *in vitro* selected ribozymes 27.3t and 27.6t^21^ were chosen for their high turnover compared to ancestors from earlier selection rounds.^20,21^ Although these ribozymes share similar secondary structures, 27.6t has an extended stem absent from the 27.3t ribozyme. PAGE analysis showed that cross-chiral ligation reactions containing the L-RNA fragments, splints, and either 27.3t or 27.6t ribozyme produced successive ligation of Fragments 2 and 3 onto Fragment 1 over time. This indicated successful assembly of L-glyS, the longest L-RNA ligated to date.

At 1 h, the 27.3t ribozyme converted 37-45% of Fragment 1 to full-length L-glyS, greatly outperforming 27.6t with only 18-25% conversion (Figure 1B). Increasing the ribozyme or L-RNA concentration showed minimal effects on ligation efficiency (Figures 1B and 1C). Additionally, L-glyS yield was typically higher after 1 h incubation compared to 16 h. This is likely due to the high magnesium levels (250 mM) required for ribozyme ligation activity,^20,21^ which can lead to RNA hydrolysis after long incubation periods. Additional 2 h and 4 h time points were tested to balance ligation progression with minimizing degradation, with 2 h found to be the optimal incubation time on average at 38-40% conversion to L-glyS (Figure 1C).

Beyond optimizing the incubation time and RNA concentrations, previous work has shown that the macromolecular crowding agent PEG-8000 can improve RNA ligation yields.^14,27,28^ By volume exclusion and altered solvent properties, PEG-8000 can increase the effective RNA concentration and promote productive intermolecular collisions. Indeed, full-length L-glyS yield increased with PEG-8000 concentration, reaching a maximum of 46% conversion at 5% PEG-8000 (Figure 2A). At higher concentrations, visible precipitation occurred along with and a drop in yield, falling to 34% conversion with 11.25% PEG-8000. Thus, 5% PEG-8000 was chosen as the optimal condition.

**Figure 2.**
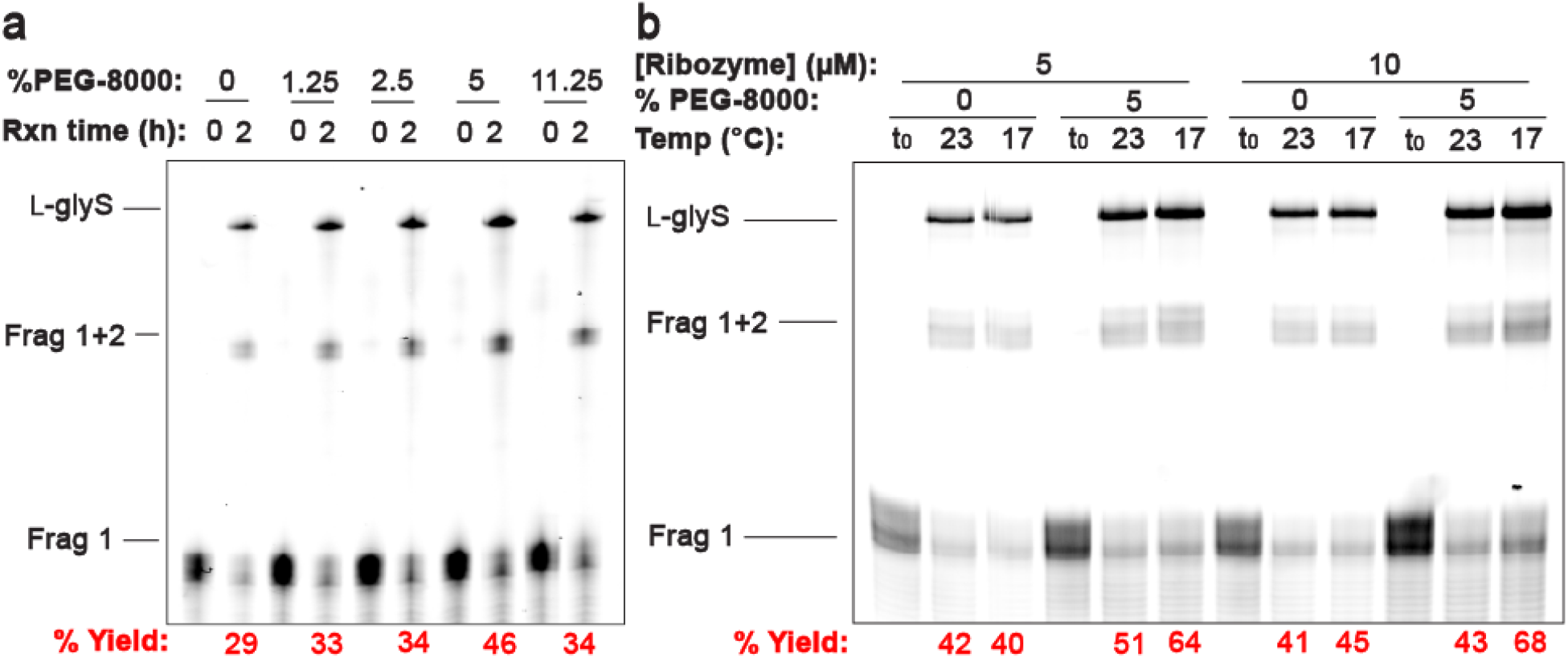
Optimization of Cross-Chiral Ligations. Denaturing PAGE analysis of cross-chiral ligations with varying concentrations of PEG-8000 incubated at **(A)** 23°C or **(B)** different temperatures. All the above reactions contained 1 µM of each splint and fragment, 5 µM of 27.3t ribozyme, 250 mM NaCl, 250 mM MgCl_2_, and 50 mM Tris (pH 8.5), and were incubated for 0 or 2 h. The label t0 defines reactions that were quenched at the 0 h time point.

Next, different reaction temperatures were investigated. Without PEG-8000, lowering the reaction temperature from 23°C to 17°C showed minimal effects on ligation efficiency (Figure 2B). However, in the presence of 5% PEG-8000, the lower temperature increased conversion to L-glyS from 43% to 68%, highlighting a synergistic effect between molecular crowding and lower temperature. This enhancement is consistent with previous reports of higher turnover of current cross-chiral ribozymes at lower temperatures,^22^ which may reflect stabilization of the ribozyme-substrate complex, improved ribozyme folding, or a combination of the two effects.

Finally, a more recently developed and higher turnover cross-chiral ligase variant 47.11 was investigated.^22^ This ribozyme previously displayed 13.6-fold faster catalytic turnover than 27.3t, enabling rapid self-amplification of its own enantiomer. However, for ligating L-glyS, 27.3t was more efficient than 47.11 (Figure S3) and was therefore used for subsequent ligation reactions.

### Comparison of *In Vitro* Performance of the L-RNA and D-RNA Glycine Sensor

For optimization and comparison with L-glyS, the original D-glyS was synthesized by *in vitro* transcription. Upon binding glycine, the fluorescence signal of glyS^8^ is activated by folding the Broccoli aptamer,^15^ which in turn binds the green-emitting fluorogenic dyes DFHBI,^29^ DFHBI-1T,^15^ and BI^16^ (Figure 3A). These dyes were analyzed with D-glyS for the best fluorescence response to glycine. With D-glyS, BI resulted in a 10-fold fluorescence enhancement response to 10 mM glycine compared to DFHBI-1T and DFHBI, which showed 7.3 and 6.0-fold turn-on, respectively (Figure 3B). Since BI was also the brightest of the dyes, it was used for subsequent assays.

**Figure 3.**
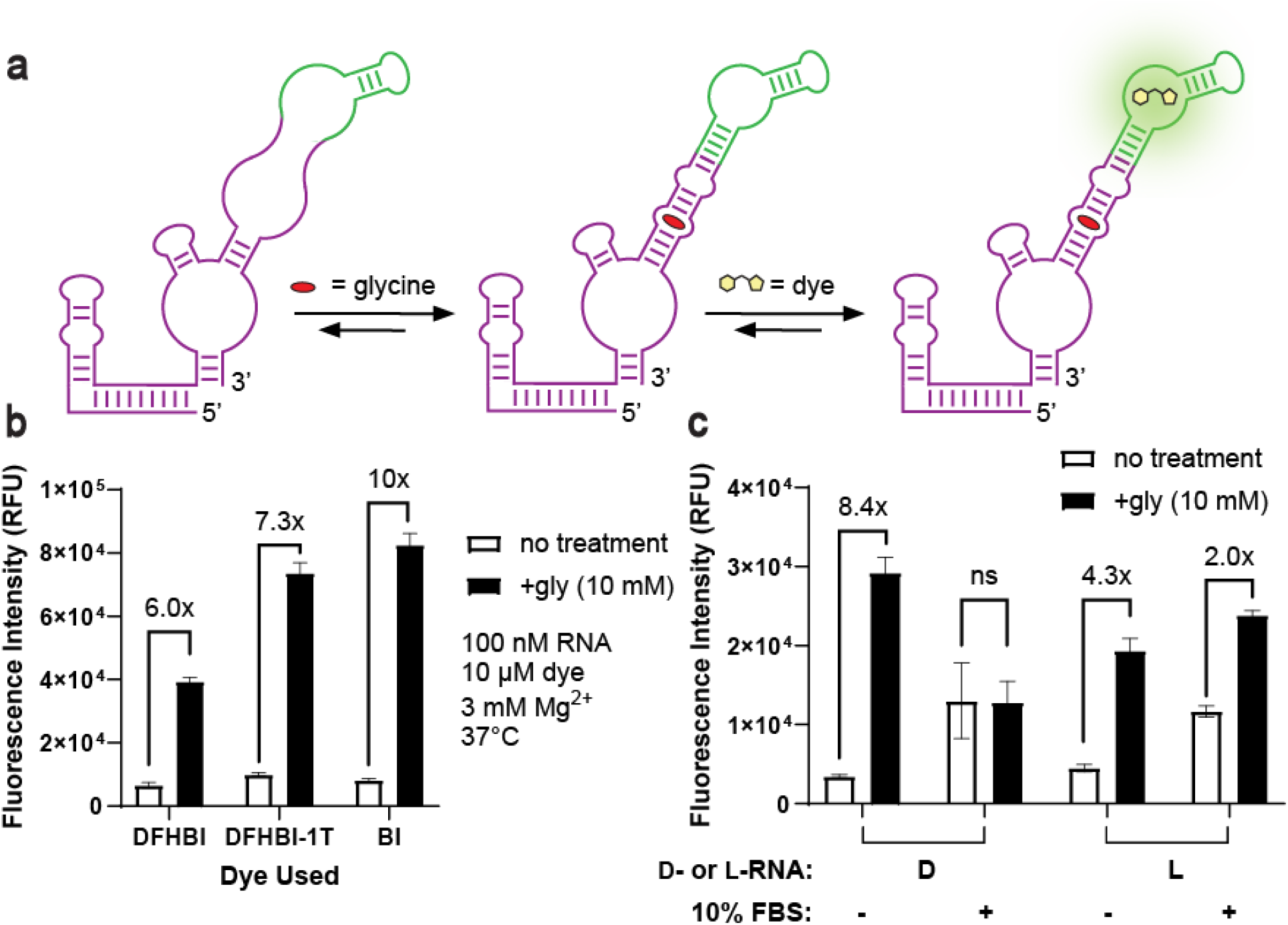
*In Vitro* Fluorescence Assays with D- and L-glyS. **(A)** Schematic of glycine sensor fluorogenic mechanism. **(B)** *In vitro* fluorescence of D-glyS with various dyes (30 µL reactions), with or without 10 mM glycine. **(C)** *In vitro* fluorescence of D- or L-glyS with 10 µM BI, with or without 10 mM glycine and 10% FBS (15 µL reactions). For all experiments, the reaction components were incubated for 30 min before measurement. The mean and standard deviation from three technical replicates are shown.

To determine which sensor performed best in complex biological environments, both D- and L-glyS were evaluated in the absence or presence of 10% fetal bovine serum (FBS) (Figure 3C). Without FBS, D- and L-glyS both showed a robust fluorescence enhancement response to 10 mM glycine. Aside from an additional 5’-G in D-glyS from IVT, the RNA sequence was the same between the two sensors. It is possible that the lower activation of L-glyS (4.3-fold) versus L-glyS (8.4-fold) may result from the additional terminal biotin and sulfo-Cy5 groups present on L-glyS but absent from D-glyS, but these groups were spaced from the L-RNA by long hexaethyleneglycol groups to avoid misfolding (Figure S1B). More likely, the discrepancy in function between D- and L-glyS may reflect the higher quality of enzymatically prepared D-RNA compared to synthetically prepared L-RNA, as observed in previous work with RNA-binding DNA aptamers.^30,31^

However, with 10% FBS present, D-glyS showed a complete loss of function, while L-glyS produced a 2-fold fluorescence enhancement response to glycine (Figure 3C). This is consistent with previous observations that fluorogenic D-RNA probes degrade within 5 minutes of adding 10% FBS compared to long-term (>12 h) nuclease stability of L-RNA.^32^ Higher background fluorescence caused the lower fold activation of L-glyS with added FBS compared to no FBS. While nuclease resistance is a primary requirement for extracellular RNA-based sensors, additional factors such as background fluorescence, sensor localization, and dye-environment interactions will also impact performance in complex biological environments. Interestingly, the background fluorescence of 10% FBS and BI together is higher than the sum of each individual component (Figure S4). This may reflect the increased viscosity and/or nonspecific protein interactions within serum, which can activate fluorogenic molecular rotor dyes like BI by hindering free intramolecular rotation and instead favoring energy release through fluorescence.^33^

Encouraged by the serum stability of L-glyS from the first ligation batch, the same ligation process was scaled up in 16 separate reactions using the remaining L-RNA fragments from SPOS. The ligation reactions were separately PAGE-purified, extracted, ethanol precipitated, then combined into one tube to generate a scaled up, concentrated batch of L-glyS. Though functional, L-glyS from the scale-up batch was less responsive than L-glyS from the initial batch (Figure S5A), despite using identical refolding conditions, appearing identical in size and integrity when measured by microfluidic electrophoresis (Figure S5B), and even after additional clean-up steps to remove leftover salts or small molecule contaminants by size exclusion spin columns. While this batch-to-batch variability hindered our ability to measure extracellular glycine, from a single solid-phase synthesis of each fragment and splint at 1 μmol scale, a total of 2.3 nmol full-length L-RNA glyS was generated, which could enable >20 cellular imaging experiments on commonly used 8-well glass slides.

## DISCUSSION

This work achieves two complementary objectives. First, cross-chiral ligation provides a route to assembling long, structurally complex L-RNAs that are difficult to access using conventional synthetic methods. Second, L-RNA enables an RBF glycine sensor to function in RNase-rich environments without rapid degradation. Previously, only cross-chiral ribozymes or short RNAs have been assembled by cross-chiral ligation,^20–22^ leaving other functional RNAs to be explored. Even though the secondary structure of glyS is highly complex, careful choice of ligation junctions ensured efficient splint annealing and ligation. This may inspire the generation of additional mirror-image riboswitch or aptamer-based sensors. Previously, the molecular crowding agent PEG-8000 has increased *in vitro* transcription yields by up to 50%.^34,35^ For RNA ligations, PEG-8000^14,27,28^ and lower reaction temperatures^22,28^ have both improved reaction efficiency. By leveraging the synergistic effect between PEG-8000 and a lower ligation temperature, up to 68% conversion to full-length L-glyS was made possible.

L-glyS, bearing additional chemical modifications relative to D-glyS, showed a lower fluorescence activation response to glycine in buffer, and there was batch-to-batch variability in L-glyS function. However, in RNase-rich serum, L-glyS outperformed D-glyS, which showed a total loss of function. This matches observations of serum tolerance with previous L-RNA-based sensors.^13,14,32^ With the current glyS designs, there is a tradeoff between maximum activation and RNase tolerance. Future improvements in L-RNA sensor design may further increase activation while maintaining RNase resistance.

While future work can improve L-glyS, the current study demonstrates an RNA-based glycine sensor that can operate in biological environments where D-RNA fails. The application of L-RNA for RNase resistance when monitoring achiral targets avoids reselection that is necessary when chemical modifications to the sugar-phosphate backbone are made due to tertiary structure differences.^11^ Although a protein-based sensor exists for imaging extracellular glycine,^25^ advantages of RNA-based glycine sensors for extracellular applications include their smaller size and that their dynamic range can be matched to varying physiological regimes using different riboswitch domains or magnesium rather than trial-and-error mutagenesis. Together, these results demonstrate that cross-chiral ligation can generate long, functional mirror-image RNAs beyond ribozymes, establish a strategic route to nuclease-resistant RNA sensors without reselection or redesign, and broaden the potential scope of glycine sensors.

## AUTHOR CONTRIBUTIONS

M.R.B. and X.H. conducted the research. M.R.B., X.H., J.T.S. and M.C.H. designed the experiments. M.R.B. and M.C.H. wrote the manuscript. J.T.S. and M.C.H. supervised the project. All authors reviewed and edited the manuscript.

## NOTES

The authors declare no competing financial interests.

## ACKNOWLEDGEMENTS

We thank Hammond and Sczepanski lab members for their helpful suggestions and advice. We thank Dr. Mike Hanson, Director of the University of Utah DNA/Peptide Synthesis Core Facility, for his assistance with solid-phase L-RNA synthesis. We thank Tessa Kilberg and Dr. Jason Shepherd for research materials and consultation on cellular experiments. This work was supported by the National Science Foundation [1815508 to M.C.H. and GRFP 2139322 to M.R.B.] and the National Institutes of Health [R01 GM124589 to M.C.H., R35GM124974 to J.T.S. and M.R.B. was supported partially by T32 GM122740]. The content is solely the responsibility of the authors and does not necessarily represent the official views of the National Institutes of Health.

