## supplemental data for "Glycine detection with a nuclease-stable L-RNA sensor"

### for

**Table of Contents** **Pages**

Materials and Methods 2 – 5

Supplementary Table 1 6

Supplementary Figures S1-S5 7 – 11

Supplementary References 12

#### Materials and Methods

##### Reagents and Oligonucleotides

All DNA oligonucleotides for RNA constructs or cloning were purchased from the University of Utah HSC Core Facility. L-ribonucleoside TBDMS phosphoramidites and L-RNA CPG solid supports were purchased from ChemGenes (Wilmington, MA). The 3'-biotinTEG CPG, Spacer phosphoramidite 18, and the 5'-Amino-Modifier C6-PDA phosphoramidite were purchased from Glen Research (Sterling, VA). Sulfo-Cy5-N-Hydroxysuccinimide (NHS) ester was purchased from Lumiprobe (Westminster, MD). DFHBI was synthesized following previously described protocols[1,2] and stored at −20°C in DMSO. DFHBI-1T was purchased from Sigma–Aldrich (St. Louis, MO) and stored at −20°C in DMSO. BI was purchased from Lucerna Technologies (Brooklyn, NY) and stored at −20°C in DMSO. All other reagents were purchased from Sigma-Aldrich (St. Louis, MO).

##### L-Oligonucleotide Synthesis

L-oligonucleotide fragments and splints were synthesized on the Applied Biosystems DNA/RNA Synthesizer 394 or the Expedite 8909 DNA Synthesizer. Fragment 1 (75 nt) was synthesized using a 2000 Å CPG, while the rest of the fragments and splints were synthesized with 1000 Å CPGs. A terminal 5'-amine was introduced to Fragment 1 using the 5'-Amino-Modifier C6-PDA phosphoramidite (Figure S1B). A 3'-biotinTEG CPG introduced a 3'-terminal biotin to Fragment 3 (Figure S1B). Terminal chemical groups were spaced from the L-RNA sequences by hexaethyleneglycol using Spacer phosphoramidite 18 (Figure S1B). All L-RNA oligos were purified by 20% denaturing PAGE before use. Excised L-RNAs were eluted overnight at room temperature in buffer with 10 mM Tris (pH 7.6), 200 mM NaCl, and 10 mM EDTA and concentrated using 3 kDa pore size Amicon Ultra Centrifugal Filter column (MilliporeSigma, Burlington, MA). Oligos were ethanol precipitated, dried, and stored in double-distilled water. The exact mass of all oligonucleotides was confirmed using the Thermo Scientific Q Exactive Focus ESI mass spectrometer, followed by deconvolution in UniDec 7.0.2.

Sulfo-Cy5 was installed onto the 5'-amine group of Fragment 1 via sulfo-Cy5 NHS ester post synthesis as described previously.[3] Briefly, the amine-bearing L-RNA (100 μM) was incubated with NHS-Sulfo-Cy5 (5 mM) in 0.1 M sodium bicarbonate (pH 8.5) with agitation on the vortexer for 16 h. The L-RNA was then precipitated with ethanol and purified by 20% denaturing PAGE.

##### Synthesis of Adenosine 5′-Diphosphate (ADP) Phosphoimidazolide (ImppA)

Adenosine 5'-diphosphate phosphoimidazolide (ImppA) was synthesized as described previously.[4] The ADP tributylaminium salt was prepared prior to synthesis to improve solubility. In short, an amberchrom 50WX8 column was equilibrated with double distilled water, followed by 70 mL of 1 M pyridine in double distilled water. To the column, a solution containing 125 mg (<0.25 mmol) adenosine 5'-diphosphate monopotassium salt in 3 mL of double distilled water was applied then eluted with 70 mL of 1:1 methanol: water mixture into 3.8 mL tributylamine in 10 mL methanol stirred on ice, at which point a white emulsion formed. This emulsion was dried using a rotary evaporator and reevaporated twice with methanol (50 mL) then once with DMF (30 mL).

Resulted yellow oil was dissolved in anhydrous DMF (7.5 mL) and stored under argon for downstream use.

Next, a separate oven-dried round-bottom flask was cooled to room temperature under argon. The cooled flask was charged with 131 mg (0.5 mmol) triphenylphosphine, 85 mg (1.25 mmol) imidazole, and 0.45 mL (1.25 mmol) triethylamine stirred in 6 mL of anhydrous DMF under argon. A solution containing 110 mg (0.5 mmol) 2,2'-dipyridyldisulfide in 1.5 mL of dry DMF was added dropwise into the stirring flask under argon, at which point the solution color was yellow-green. After which, previously prepared ADP tributylamine salt in 7.5 mL DMF was added dropwise with stirring under argon, and the solution color changed from yellow-green to warm yellow. The reaction was allowed to stir for an additional 3 h under argon.

The reaction was added dropwise to an open-air stirring solution containing 0.55g (4.5 mmol) sodium perchlorate, 55 mL acetone, and 27.5 mL anhydrous diethyl ether. The precipitant was collected via centrifugation for 5 min at 5000 rpm, then washed twice with acetone followed by diethyl ether. The crude product was dried under vacuum, characterized via ESI-MS, then used without further purification.

##### Adenosyl-Triphosphorylation of L-RNAs

In a microcentrifuge tube, 5 nmol of dried 5'-phosphorylated Fragment 2 or 3 was resuspended in a 100 µL solution containing 100 mM ImppA and 50 mM $MgCl_2$. The mixture was allowed to incubate at 52.6 °C for 3 hours, with an additional freshly prepared 50 µL of 100 mM ImppA in 50 mM $MgCl_2$ added halfway. Reaction mixture was quenched, desalted, and purified by 20% denaturing PAGE as described earlier. The masses of the adenosyl-triphosphorylated L-RNA oligos were verified by mass spectrometry.

##### *In Vitro* Transcription of D-RNA Constructs

For *in vitro* transcription, DNA templates were amplified with Phusion DNA Polymerase (UC Berkeley MacroLab) or Q5 DNA Polymerase (New England Biolabs) using sequence-confirmed plasmids as templates. The T7 promoter was contained within the template or appended via an overhang on 5′ end of the forward primer. The PCR product was purified and concentrated using the QIAquick PCR Purification Kit (Qiagen).

RNA was transcribed from the purified PCR product as previously described.[5] Briefly, ~1 µg of DNA was incubated at 37°C for 4 h with homemade T7 RNA polymerase in 40 mM Tris–HCl, pH 8.0, 6 mM $MgCl_2$, 2 mM spermidine, 10 mM DTT, 1 U of inorganic pyrophosphate, and 2 mM rNTPs. The RNAs were purified by denaturing (7.5 M urea) 6% polyacrylamide gel electrophoresis (PAGE). PAGE-purified RNA was visualized by UV shadowing and eluted from gel pieces using a solution of 10 mM Tris–HCl, pH 7.5, 200 mM NaCl, and 1 mM EDTA, pH 8.0. Extracted RNAs were ethanol precipitated, dried, and resuspended in $ddH_2O$. Glycogen was omitted when precipitating cross-chiral ribozymes, so that the amount of crowding agent in cross-chiral ligations could be tightly controlled. To accurately quantify the RNA, a neutral pH thermal hydrolysis assay was used to remove the hypochromic effect from RNA secondary structure.[6]

**Small-Scale Cross-Chiral Ligation Reactions to Synthesize L-RNA Glycine Sensor**

*Initial Cross-Chiral Ligations*

The L-RNA splints, L-RNA fragments, (final concentrations: 1 µM each), and the D-RNA 27.3t or 27.6t cross-chiral ligase ribozyme (final concentration: 5 or 10 µM) were combined in a reaction mixture (5 µL) containing 250 mM NaCl and 50 mM Tris (pH 8.5). This mixture was heated to 70°C for 2 min, then slowly cooled to 20 °C at a ramp rate of 0.1 °C/s. The reaction was initiated by addition of 5 µL of 500 mM $MgCl_2$, 250 mM NaCl, and 50 mM Tris (pH 8.5). The reaction was incubated at 23 °C for 0, 1, or 16 h. For analyzing reaction progression, 2 µL aliquots were quenched with 18 µL of 90% v/v formamide and 50 mM EDTA. Then, 10 µL of each of these mixtures was analyzed by 12% denaturing PAGE. The gels were imaged on the Typhoon with the Cy5 filter set and quantified using ImageJ.

*Determining an Optimal Temperature*

The L-RNA splints, L-RNA fragments, (final concentrations: 1 µM each), D-RNA 27.3t cross-chiral ligase ribozyme (final concentration: 5 or 10 µM), and PEG-8000 (final concentration: 0 or 5%) were combined in a reaction mixture (5 µL) containing 250 mM NaCl and 50 mM Tris (pH 8.5). This mixture was heated to 70°C for 2 min, then slowly cooled to 20 °C at a ramp rate of 0.1 °C/s. The reaction was initiated by addition of 5 µL of 500 mM $MgCl_2$, 250 mM NaCl, and 50 mM Tris (pH 8.5). The reaction was incubated at 23 °C or 17 °C for 0 or 2 h. For analyzing reaction progression, 2 µL aliquots were quenched with 18 µL of 90% v/v formamide and 50 mM EDTA. Then, 10 µL of each of these mixtures was analyzed by 12% denaturing PAGE. The gels were imaged on the Typhoon with the Cy5 filter set and quantified using ImageJ.

*Determining an Optimal Incubation Time*

The L-RNA splints, L-RNA fragments, (final concentrations: 1, 1.5, or 2 µM each), and D-RNA 27.3t cross-chiral ligase ribozyme (final concentration: 5 µM) were combined in a reaction mixture (5 µL) containing 250 mM NaCl and 50 mM Tris (pH 8.5). This mixture was heated to 70°C for 2 min, then slowly cooled to 20 °C at a ramp rate of 0.1 °C/s. The reaction was initiated by addition of 5 µL of 500 mM $MgCl_2$, 250 mM NaCl, and 50 mM Tris (pH 8.5). The reaction was incubated at 23 °C for 0, 1, 2, or 4 h. For analyzing reaction progression, 2 µL aliquots were quenched with 18 µL of 90% v/v formamide and 50 mM EDTA. Then, 10 µL of each of these mixtures was analyzed by 12% denaturing PAGE. The gels were imaged on the Typhoon with the Cy5 filter set and quantified using ImageJ.

*Determining an Optimal PEG-8000 Concentration*

The L-RNA splints, L-RNA fragments, (final concentrations: 1 µM each), D-RNA 27.3t cross-chiral ligase ribozyme (final concentration: 5 µM), and PEG-8000 (final concentration: 0, 1.25, 2.5, 5, or 11.25%) were combined in a reaction mixture (5 µL) containing 250 mM NaCl and 50 mM Tris (pH 8.5). This mixture was heated to 70°C for 2 min, then slowly cooled to 20 °C at a ramp rate of 0.1 °C/s. The reaction was initiated by addition of 5 µL of 500 mM $MgCl_2$, 250 mM NaCl, and 50 mM Tris (pH 8.5). The reaction was incubated at 23 °C for 0 or 2 h. For analyzing reaction progression, 2 µL aliquots were quenched with 18 µL of 90% v/v formamide and 50 mM EDTA. Then, 10 µL of each of these mixtures was analyzed by 12% denaturing PAGE. The gels were imaged on the Typhoon with the Cy5 filter set and quantified using ImageJ.

*Determining an Optimal Ribozyme*

The L-RNA splints, L-RNA fragments, (final concentrations: 1 μM each), D-RNA 27.3t or 47.11 cross-chiral ligase ribozyme (final concentration: 5 μM), and PEG-8000 (final concentration: 0 or 5%) were combined in a reaction mixture (5 μL) containing 250 mM NaCl and 50 mM Tris (pH 8.5). This mixture was heated to 70°C for 2 min, then slowly cooled to 20 °C at a ramp rate of 0.1 °C/s. The reaction was initiated by addition of 5 μL of 500 mM $MgCl_2$, 250 mM NaCl, and 50 mM Tris (pH 8.5). The reaction was incubated at 23 °C for 0 or 2 h. For analyzing reaction progression, 2 μL aliquots were quenched with 18 μL of 90% v/v formamide and 50 mM EDTA. Then, 10 μL of each of these mixtures was analyzed by 12% denaturing PAGE. The gels were imaged on the Typhoon with the Cy5 filter set and quantified using ImageJ.

##### Scale-Up Cross-Chiral Ligation Reactions to Synthesize L-RNA Glycine Sensor

The L-RNA splints and L-RNA fragments (final concentration: 1 μM each), D-RNA cross-chiral ligase ribozyme 27.3t (final concentration: 10 μM), and PEG-8000 (final concentration: 5%) were combined in a reaction mixture (25 μL) containing 250 mM NaCl and 50 mM Tris (pH 8.5). This mixture was heated to 70°C for 2 min, then slowly cooled to 20 °C at a ramp rate of 0.1 °C/s. The reaction was initiated by addition of 25 μL of 500 mM $MgCl_2$, 250 mM NaCl, and 50 mM Tris (pH 8.5). The reaction was incubated at 17 °C for 2 h. The RNAs were purified by denaturing (7.5 M urea) 6% polyacrylamide gel electrophoresis (PAGE). PAGE-purified RNA was visualized by UV shadowing and eluted from gel pieces using a solution of 10 mM Tris–HCl, pH 7.5, 200 mM NaCl, and 1 mM EDTA, pH 8.0. Extracted RNAs were ethanol precipitated, dried, and resuspended in $ddH_2O$. To accurately quantify the RNA, a neutral pH thermal hydrolysis assay was used to remove the hypochromic effect from RNA secondary structure.

##### *In Vitro* Fluorescence Assays

*In vitro* fluorescence assays were performed as described previously.[5] In binding buffer containing 40 mM HEPES (pH adjusted to 7.5 with KOH), 125 mM KCl, 10 μM dye, and varying concentrations of $MgCl_2$, RNA, and glycine. Before all assays, RNA (10×) was renatured by heating at 70°C for 3 min followed by 15 min of slowly cooling to room temperature in binding buffer. For the fluorescence assays, the dye (DFHBI, DFHBI-1T, or BI) was added to the reaction mixture in a Greiner Bio-One 384-well black plate containing binding buffer, RNA, glycine (as indicated), and FBS (as indicated) for a total reaction volume of 15-30 μL as indicated. After incubation in the plate for 30 min at 37 °C, the fluorescence emission was measured by the SpectraMax i3x plate reader (Molecular Devices) at the dye's excitation/emission maxima and averaged across technical replicates. The fluorescence turn-on of glycine sensors was calculated by dividing fluorescence in the presence of glycine by fluorescence in the absence of glycine.

**Supplementary Table 1.** RNA sequences used in cross-chiral work

| Name | D- or L-RNA | Sequence (5' to 3') |
|---|---|---|
| Fragment 1 | L | **Amino**-**Sp18**-CCGAAUGAAGUUUAAUAGAAGUGGGGAAUCGUUUGAU UUUCCAUGACUGUAAAUGGACGGAACUCUGGAGAGACC |
| Fragment 2 | L | **Phosphate**-GUAAAGGCACCGAAGGGGCAAGGCCUGUCGAGUAGAGU GUGGGCUC |
| Fragment 3 | L | **Phosphate**-GCAAGAGACGGUCGGGUCCAGGCUCAAACUCUCAGGUA AAAGGACAGAG-**Sp18**-**Biotin** |
| Splint 1 | L | CUUCGGUGCCUUUACGGUCUCUCCAGAGUU |
| Splint 2 | L | CCGACCGUCUCUUGCGAGCCCACACUCUAC |
| L-glyS | L | **Cy5**-**Sp18**-CCGAAUGAAGUUUAAUAGAAGUGGGGAAUCGUUUGAUUU UCCAUGACUGUAAAUGGACGGAACUCUGGAGAGACCGUAAAGGCACCG AAGGGGCAAGGCCUGUCGAGUAGAGUGUGGGCUCGCAAGAGACGGUC GGGUCCAGGCUCAAACUCUCAGGUAAAAGGACAGAG-**Sp18**-**Biotin** |
| D-glyS | D | GCCGAAUGAAGUUUAAUAGAAGUGGGGAAUCGUUUGAUUUUCCAUGAC UGUAAAUGGACGGAACUCUGGAGAGACCGUAAAGGCACCGAAGGGGC AAGGCCUGUCGAGUAGAGUGUGGGCUCGCAAGAGACGGUCGGGUCCA GGCUCAAACUCUCAGGUAAAAGGACAGAG |
| 27.3t | D | GGGGUGGUGGACGUGAUCAUUACGGAUCACUAACUCGUCAGUGCAUU GAGAAGGAGAAUAAAAUGCACAUAGGUCGAAAGACCUUAUACAAGAAC UGUAUCACCGGAGGGCGAGCACCACC |
| 27.6t | D | GGGGUGGCGGACGUGUUUCACGUAUAAUCGUGCGGGACACUGACUCG UCAGUGCAUUGAGAAGGAGGAUAAAAUGCACAUAGGUCGAAAGACCUU AUACAAGAACUGUAUCACCGGAGGGCGAGCACCACC |
| 47.11 | D | GGGACAUGGGUGUUGGACUUGAUCAUUACGGAUCAAUAACUCGUCAG CGCAUUCAGAAGGAGAAGAAAAUGCGCAUAGGUCAAAAGACCUUAUAC AAGAACUGUAUCCCCGGAGGGCGAGCACCACCCAU |

.

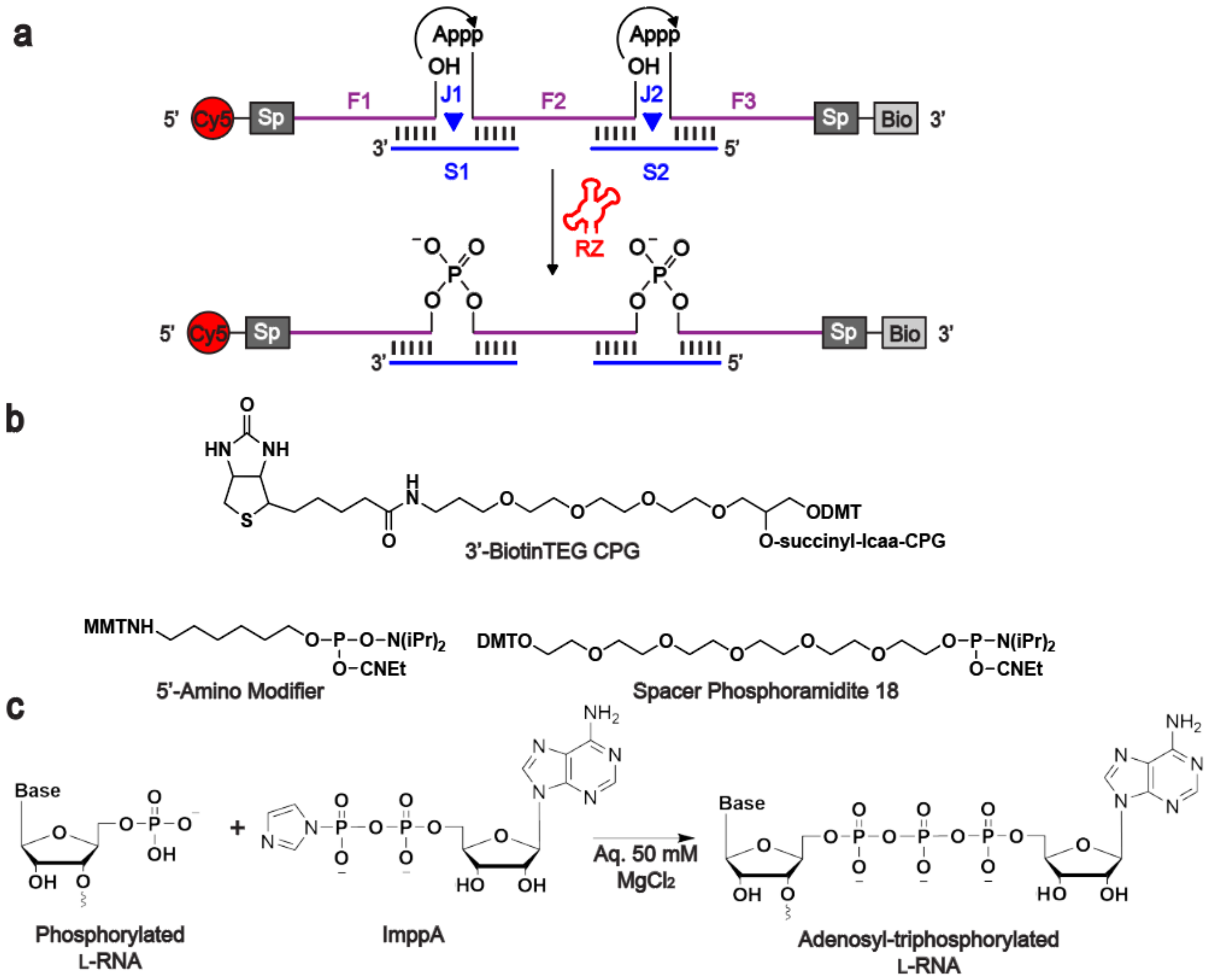


**Supplementary Figure 1. Schematic of cross-chiral ligation. (A)** In cross-chiral ligations, a D-RNA cross-chiral ribozyme (RZ) can join L-RNA glyS fragments (F1-F3) together at ligation junctions (J1, J2) when templated by L-RNA splints (S1, S2). Importantly, F2 and F3 display 5'-adenosyltriphosphate (Appp) groups. "Sp" indicates hexaethyleneglycol spacer regions, and "Bio" indicates biotin triethylene glycol. **(B)** Structures of reagents used to install amino, spacer, and biotin modifications during solid-phase L-RNA synthesis. "CNEt" indicates cyanoethyl, "DMT" indicates dimethoxytrityl, "MMT" indicates monomethoxytrityl, "iPr" indicates isopropyl, and "lcaa-CPG" indicates the long-chain alkyl amine connected to the controlled-pore glass support. **(C)** Schematic of the chemical adenosyltriphosphorylation.

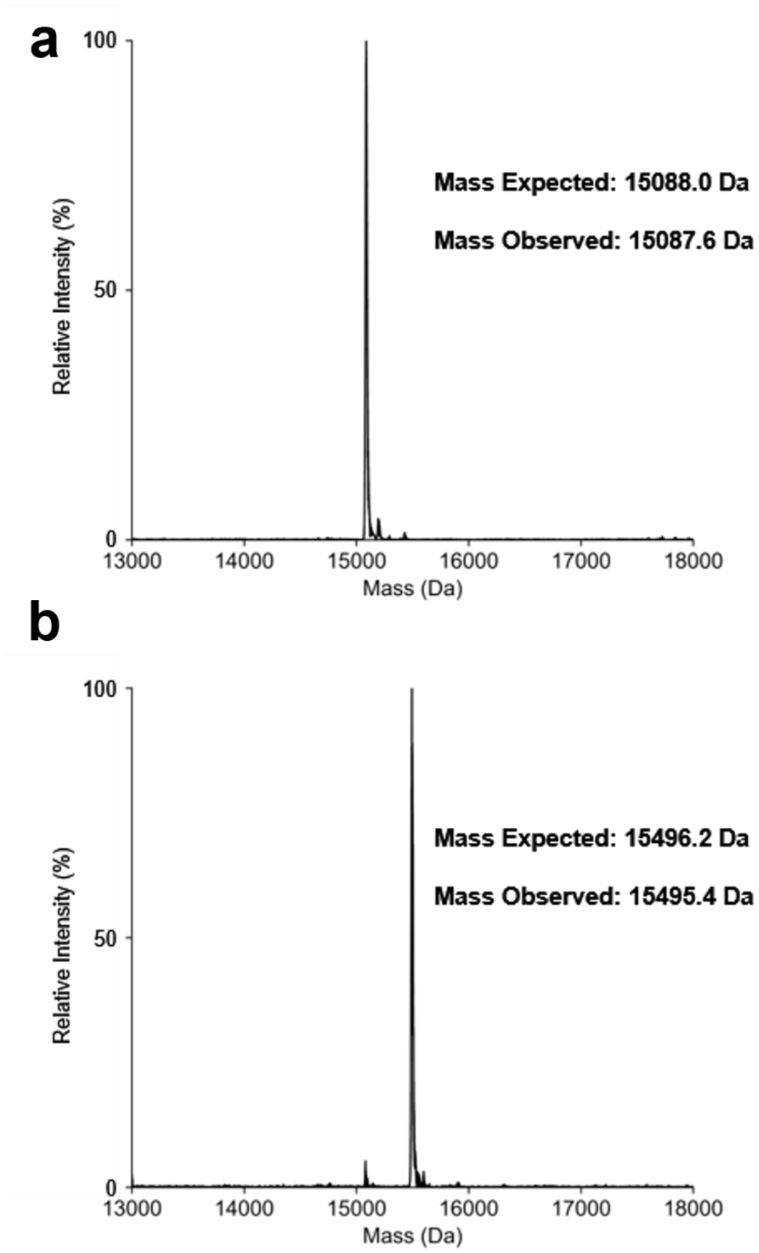


**Supplementary Figure 2. Mass spectrometry analysis of L-RNA adenosyl-triphosphorylation.** Representative mass spectrum of L-RNA Fragment 2 before **(A)** or after **(B)** adenosyl-triphosphorylation using ImppA.

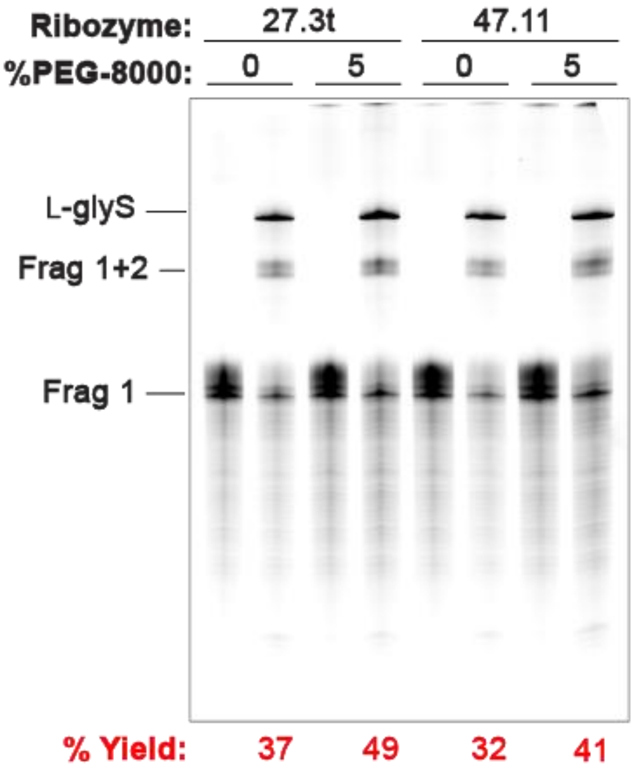


**Supplementary Figure 3. Cross-chiral ligation with various ribozymes.** Analysis of cross-chiral ligations with using 27.3t or 47.11 ribozymes and PEG-8000 concentration of 0 or 5%. All the above reactions contained 1 µM of each splint and fragment, 5 µM of ribozyme, 250 mM NaCl, 250 mM $MgCl_2$, and 50 mM Tris (pH 8.5).

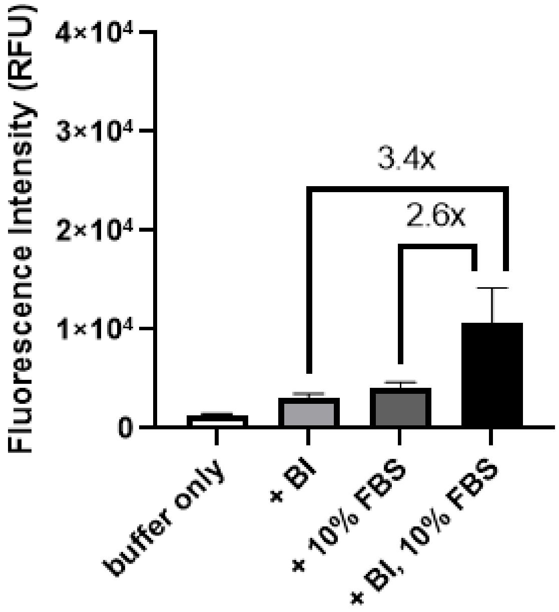


**Supplementary Figure 4. Control *in vitro* fluorescence assay without RNA.** *In vitro* fluorescence of buffer with BI, 10% FBS, or both (15 µL reactions). The reaction components were incubated for 30 min at 37°C before measurement. The mean and standard deviation from three technical replicates is shown.

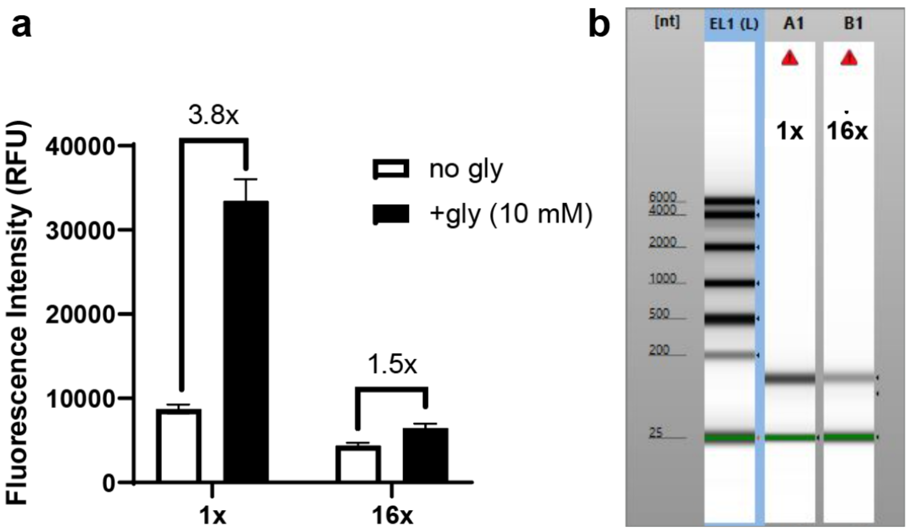


**Supplementary Figure 5. Analyzing different batches of L-glyS. (A)** *In vitro* fluorescence of 1x or 16x-scale batches of L-glyS (15 µL reactions). The reaction components were incubated for 30 min before measurement. The mean and standard deviation from two technical replicates is shown. **(B)** Microfluidic electrophoresis on the Tapestation of the 1x and 16x batches of 170-nt L-glyS. EL1 (L) is the electronic ladder used to estimate the length of the RNA.
